# The impact of FASTQ and alignment read order on structural variation calling from long-read sequencing data

**DOI:** 10.1101/2023.03.27.534439

**Authors:** Kyle Lesack, James D. Wasmuth

## Abstract

**Background:** Structural variation (SV) calling from DNA sequencing data has been challenging due to several factors, such as the ambiguity of short-read alignments, multiple complex SVs in the same genomic region, and the lack of “truth” datasets for benchmarking. Additionally, caller choice, parameter settings, and alignment method are known to affect SV calling. However, the impact of FASTQ read order on SV calling has not been explored for long-read data.

**Results:** In this study, we used PacBio DNA sequencing data from 15 *Caenorhabditis elegans* isolates to evaluate the dependence of different SV callers on FASTQ read order. Comparisons of variant call format (VCF) files generated from the original and permutated FASTQ files demonstrated that the order of input data had a large impact on SV prediction, particularly for pbsv. The overall differences were lowest for Sniffles, regardless of the aligner used. The type of variant most affected by read order varied by caller. For pbsv, most differences occurred for deletions and duplications, while for Sniffles, permutating the read order had a stronger impact on insertions. For SVIM, inversions and deletions accounted for most differences.

**Conclusion:** The results of this study highlight the dependence of SV calling on the order of reads encoded in FASTQ files, which has not been recognized in long-read approaches. These findings have implications for the replication of SV studies and the development of consistent SV calling protocols. Our study suggests that researchers should pay attention to the order of reads when analyzing long-read sequencing data for SV calling.

## Introduction

Structural variants (SVs) describes a broad category of large—typically ≥ 100bp—chromosome alterations that play key evolutionary roles as a source of novel genes and adaptive phenotypes^1^. Common variants include deletions, insertions, inversions, translocations, and duplications.

The detection of SVs from DNA sequencing data has proved to be a challenging and the barriers to accurate and comprehensive SV prediction are multifactorial. Many variants span genomic regions that exceed the read sizes generated by high-throughput sequencing platforms, which may hinder accurate mapping to a reference genome^2^. Because SVs, such as deletions and duplications, typically arise in low complexity regions, this problem is compounded by ambiguous mappings that are inherent to short-read alignments^3^. Multiple SVs may also occur in the same genomic region^4–6^, leading to complex rearrangements that are difficult to resolve computationally^7^.

Considerable work has gone into improving SV calling methods, but the limited availability of “truth” datasets with known variants has limited benchmarking efforts^8–10^. Several sources of variability have also been identified that contribute further difficulties for the development of consistent SV calling protocols. While differences in library preparation^11^ or sequencing platform can affect the predicted SVs, considerable disparities between the call sets generated by different sequencing centers has been observed when using the same protocols^12^. On the computational side, caller choice, parameter settings, and alignment method are known to affect SV calling^13,14^. For short-read data, how software handles ambiguous read-to-genome mappings is a surprising and significant source of variation in SV identification; changing the order of the reads in the FASTQ file led to changes in predicted SVs^15^. These discrepancies were as high as 25% for certain callers, raising the possibility that the random nature of FASTQ read order could have a substantial impact on replication work. Without knowing ground truth, it is not possible to quantify the exact relationship between these SV call discrepancies and caller accuracy. Nonetheless, high dependence on read order reflects poor performance for certain metrics. Theoretically, a given caller may have a high true positive rate but may also generate considerably different predictions when the FASTQ file read order is changed. This would be consistent with the caller having high precision, but low recall, as the call set differences would include many true positives that are members of one call set but not the other. However, a caller with a high false positive rate could conceivably have a high recall, while also exhibiting high read order dependence. Here, most of the call set differences would correspond to false positives generated from a specific read order that were not generated in the other. An interesting question would be if the calls that differ following the randomization of read order are biased towards being false positives? If any biases are found to exist, read order permutation could be used to improve the caller performance.

It is unclear if FASTQ read order should also be a consideration for SV calling from long-read data. To evaluate the FASTQ read order dependence of long-read callers, we used PacBio DNA sequencing data from 15 separate *Caenorhabditis elegans* isolates. We generated FASTQ files with permutated read order from the original files and evaluated the differences between the SVs predicted using the initial and randomized data. Although each caller was found to be deterministic, the order of reads provided to each caller had an impact on the predicted SVs. Several factors were identified that contributed the dependence of different SV callers on the order of FASTQ file reads. These results bring attention to a largely unrecognized factor that may affect the inferences made from structural variation studies.

## Methods

### DNA Sequencing Datasets and Read Order Randomization

PacBio sequencing data were obtained for 14 *Caenorhabditis elegans* isolates from the *Caenorhabditis elegans* Natural Diversity Resource (CeNDR) database^16^ and as well as one sequencing run of the reference strain, N2 (SRA accession = DRR142768). For each sequencing run, we created a permutated FASTQ file with randomized orders of the original reads. Initially, the BBTools (v39.00)^17^ shuffle.sh and shuffle2.sh scripts were used to randomize the FASTQ sequence order. However, in addition to randomizing the sequence order, the FASTQ files generated by both scripts contained changes in the Phred scores. Specifically, Phred scores encoded as “!” in the original FASTQ files were changed to “#” in the permutated versions. Therefore, the permutated FASTQ files were created using the permutated seq-shuf^18^ script. A comparison between the original and permutated FASTQ files indicated that only the order of sequences was altered using seq-shuf.

### Sequence Alignment, subsampling, and Structural Variant Prediction

Alignments for each FASTQ file were created using pbmm2 (v1.7.0)^19^, Minimap2 (v2.17)^20^, and NGMLR (v0.2.7)^2^. The BAM files were then sorted using SAMtools (v1.9)^21^ and Picard Tools (v2.27.5)^22^. SVs were predicted from the original and permutated datasets using the default parameters for three tools: pbsv (v2.8.0)^23^, Sniffles (v2.0.6)^2^, and SVIM (v2.0.0)^24^. We were unable to generate SV predictions in pbsv from the Minimap2 and NGMLR alignments, therefore, the pbsv predictions were limited to its recommended aligner, pbmm2. Variant calls below 100bp were filtered out in order to limit the analysis to structural variants. We confirmed that each caller was deterministic by calling SVs twice using each alignment file.

The median sequencing depths (Minimap2 median depth = 145.602; NGMLR median depth = 135.136; pbmm2 median depth = 136.831) were higher than those often used for structural prediction studies. Therefore, each BAM file was subsampled to four separate depths (10X, 20X, 40X, and 60X) using SAMtools (v1.9)^21^ to examine the read order dependence at depths consistent with those generated from larger genomes.

### Comparison of Predicted Structural Variants

For each isolate, we compared the predictions generated from the original and permutated FASTQ files from the same aligner. Two approaches were used to assess the impact of sequence randomization on the predicted structural variants: (1) vcf-level differences and (2) coordinate-level differences. At the VCF-level, the SV calls contained in the variant call format (VCF) files created from the original and permutated FASTQ files were compared using all VCF fields, except for the variant ID. The variant ID field was excluded because it only describes an arbitrary name assigned to the SV call from its order in the VCF file. Because an analyst may only be concerned with the locations of SVs that pass quality filtering, a less strict analysis was also performed that limited comparisons to the variant type, filter, chromosome, start coordinate, and end coordinate (not applicable to INS calls).

For each comparison, the symmetric difference (the set of SVs predicted from either the original or permutated FASTQ files, but not in their intersection) was used to quantify the differences that occurred following the randomization of read order. The proportions in the text were represented using the mean ± standard deviation. The counts included in the figures indicate the mean proportion of different calls that resulted from read order permutation.

### Code Availability

Snakemake (v7.9.0)^25^ was used to manage the individual scripts used for read order permutation, read alignment, subsampling, SV prediction, and statistical summary. The entire workflow is available at https://github.com/kyleLesack/pacbio_read_order_shuffling.

## Results

### Structural Variation Calling Is Affected by FASTQ Read Order

Comparisons of VCF files generated from the FASTQ files containing reads in the original and permutated orders demonstrated that the order of input data had a large impact on SV prediction (Figure 1). While each caller exhibited dependence on FASTQ read order, the overall impact was substantially higher for pbsv, with a mean difference of 0.597 ± 0.144 (Table 1). The overall differences were lowest for Sniffles, regardless of the aligner used (Minimap2 = 0.018 ± 0.009; NGMLR = 0.023 ± 0.010; pbmm2 = 0.017 ± 0.009). The alignment method had a greater impact on SVIM, as fewer different calls were predicted from NGMLR alignments (0.048 ± 0.035) compared to those from pbmm2 (0.058 ± 0.033) and Minimap2 (0.076 ± 0.071).

**Figure 1.**
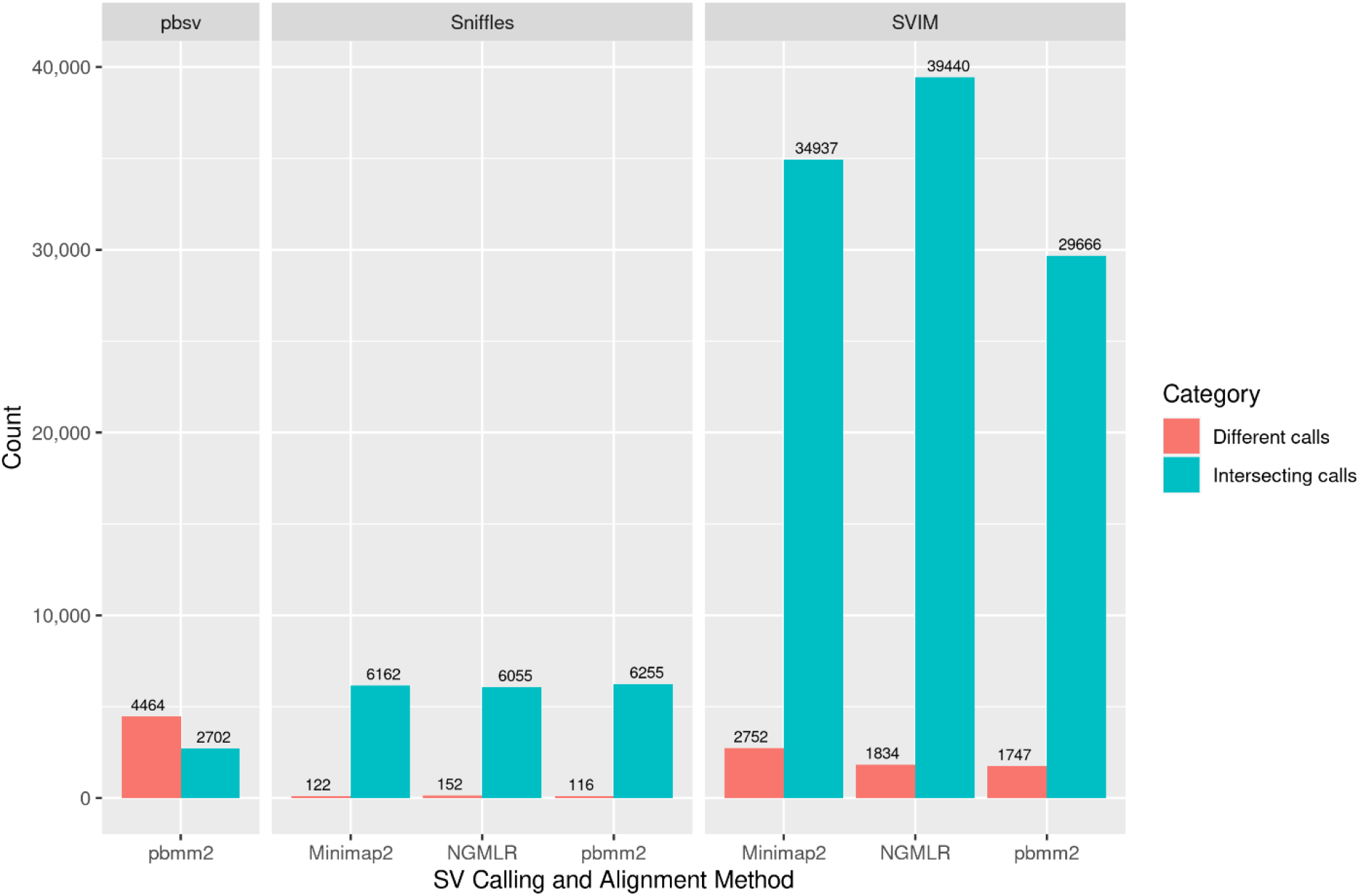
Comparison of SVs Predicted from FASTQ Files Containing the Original and Permutated Read Orders. Intersecting calls describe SVs predicted from both the original and permutated FASTQ files. Different calls include all SVs predicted exclusively from either the original or permutated FASTQ files. All VCF file fields, except for the variant ID, were required to be equal in both the original and permutated SV calls to be classified as intersecting. The counts describe the mean values for the predictions generated using the same aligner and caller.

**Table 1.**
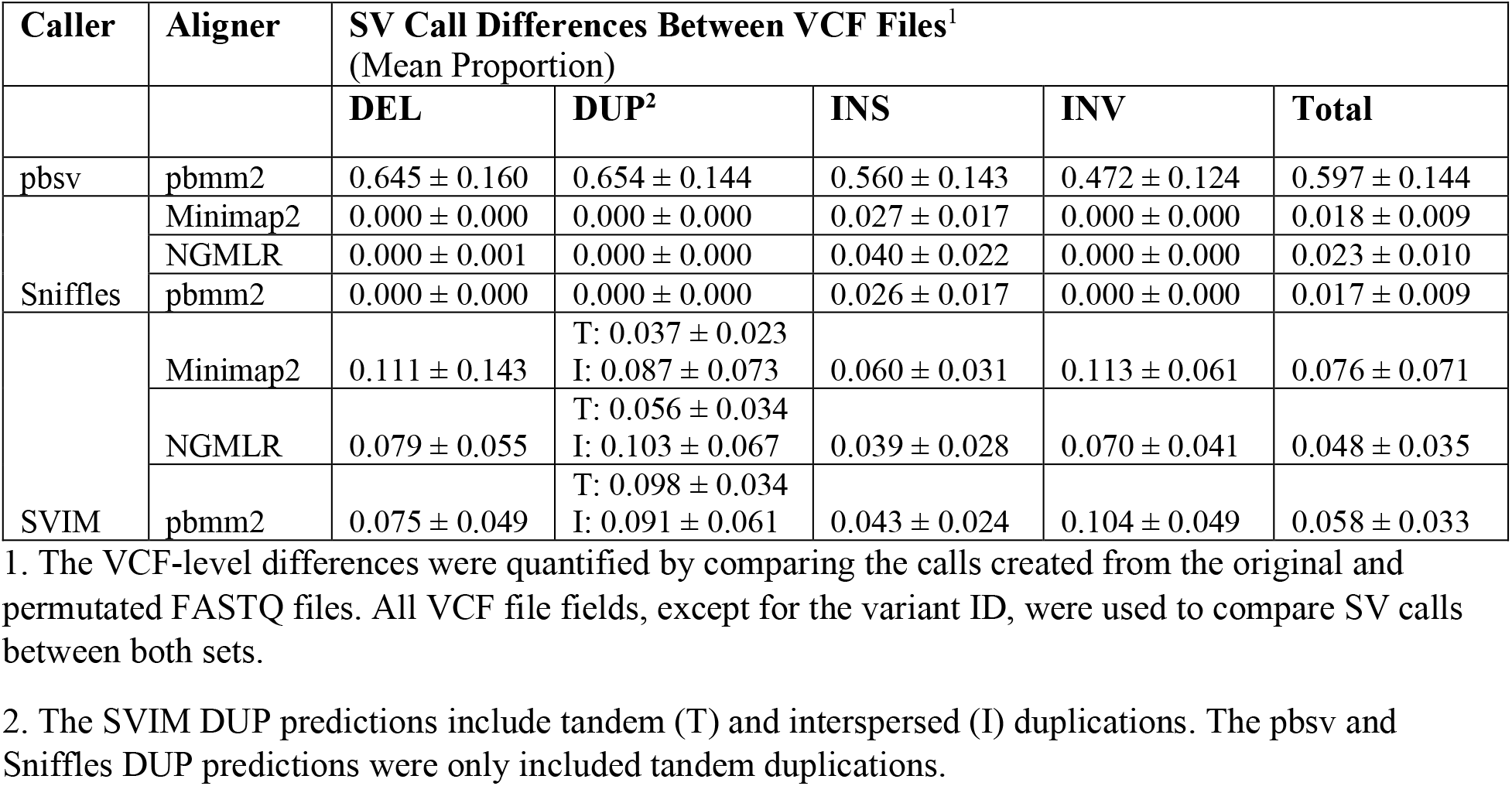
Differences in VCF File SV Calls Between the Original and Permutated Datasets.

The type of variant most affected by read order varied by caller. For pbsv, most differences occurred for deletions (0.645 ± 0.160) and duplications (0.654 ± 0.144). For Sniffles, permutating the read order had a stronger impact on insertions (Minimap2 = 0.027 ± 0.017; NGMLR = 0.040 ± 0.022; pbmm2 = 0.026 ± 0.017). For SVIM, inversions (Minimap2 = 0.113 ± 0.061; NGMLR = 0.070 ± 0.041; pbmm2 = 0.104 ± 0.049) and deletions (Minimap2 = 0.11; NGMLR = 0.08; pbmm2 = 0.07) accounted for most differences.

### BAM File Sorting Contributes to FASTQ Read Order Dependence in SV Calling

Several analyses were included to identify potential causes of the differences observed following the randomization of FASTQ file read order. Variant calling was repeated using the same alignment files, but identical results were obtained from each alignment, indicating that the callers are deterministic (i.e., the same results are obtained given identical input data). For each isolate, the alignments generated from the original and permutated FASTQ files were compared. For each read alignment in original BAM file, an identical alignment was present in the permutated version. However, despite being sorted using SAMtools, the order of aligned sequences differed between the two BAM files. Further examination revealed that the order of alignments varied for sequences aligned to the same leftmost genomic coordinate, which is also described in the SAMtools documentation (http://www.htslib.org/doc/samtools-sort.html). Identical results were obtained from the original and permutated FASTQ files when they were aligned using Picard.

### Most Differences Do Not Affect Genomic Coordinates

For each caller, fewer different calls were observed when the comparison was limited to the VCF filter category (e.g., PASS) and genomic coordinates (Figure 2). While less stringent than an exact comparison of all VCF fields, these criteria may be of interest to the analyst that does not perform further post hoc quality assurance following SV calling. Unsurprisingly, these criteria resulted in fewer call set differences for each caller (Table 2), with pbsv again being the most dependent on the FASTQ file read order (0.100 ± 0.032). For Sniffles, permutating the read had a negligible impact on the predicted SVs, as differences occurred for only seven SV calls (three of 94,536 from the pbmm2 alignments; four of 91,785 from the NGMLR alignments). For SVIM, the alignment method again affected the differences observed between the original and permutated call sets (NGMLR = 0.013 ± 0.009; pbmm2 = 0.020 ± 0.010; Minimap2 = 0.040 ± 0.010).

**Figure 2.**
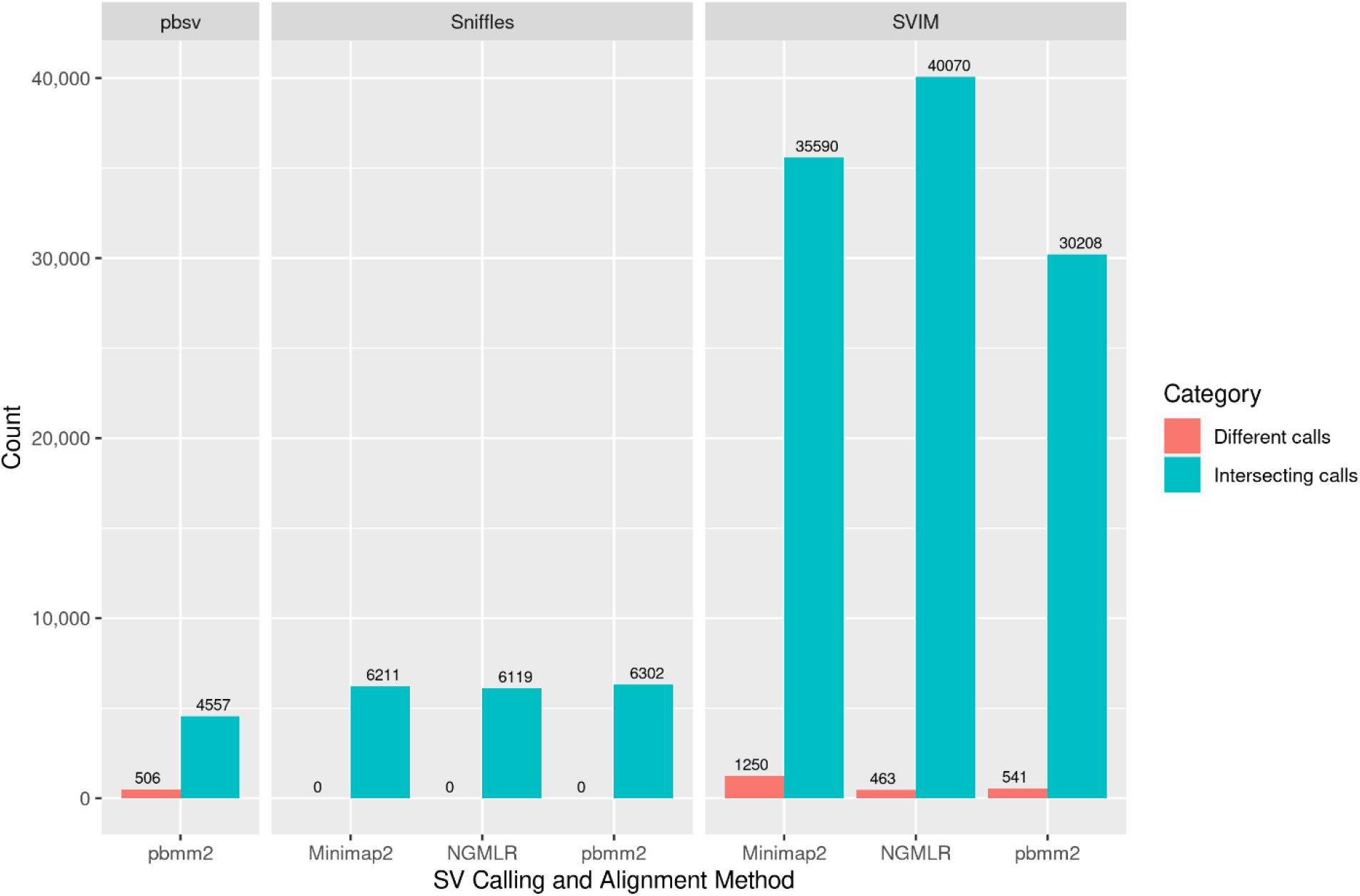
Comparison of SVs Predicted from FASTQ Files Containing the Original and Permutated Read Orders. Intersecting calls describe SVs predicted from both the original and permutated FASTQ files. Different calls include all SVs predicted exclusively from either the original or permutated FASTQ files. For each SV type, the VCF filter category and genomic coordinates were required to match to be classified as intersecting. The counts describe the mean values for the predictions generated using the same aligner and caller.

**Table 2.**
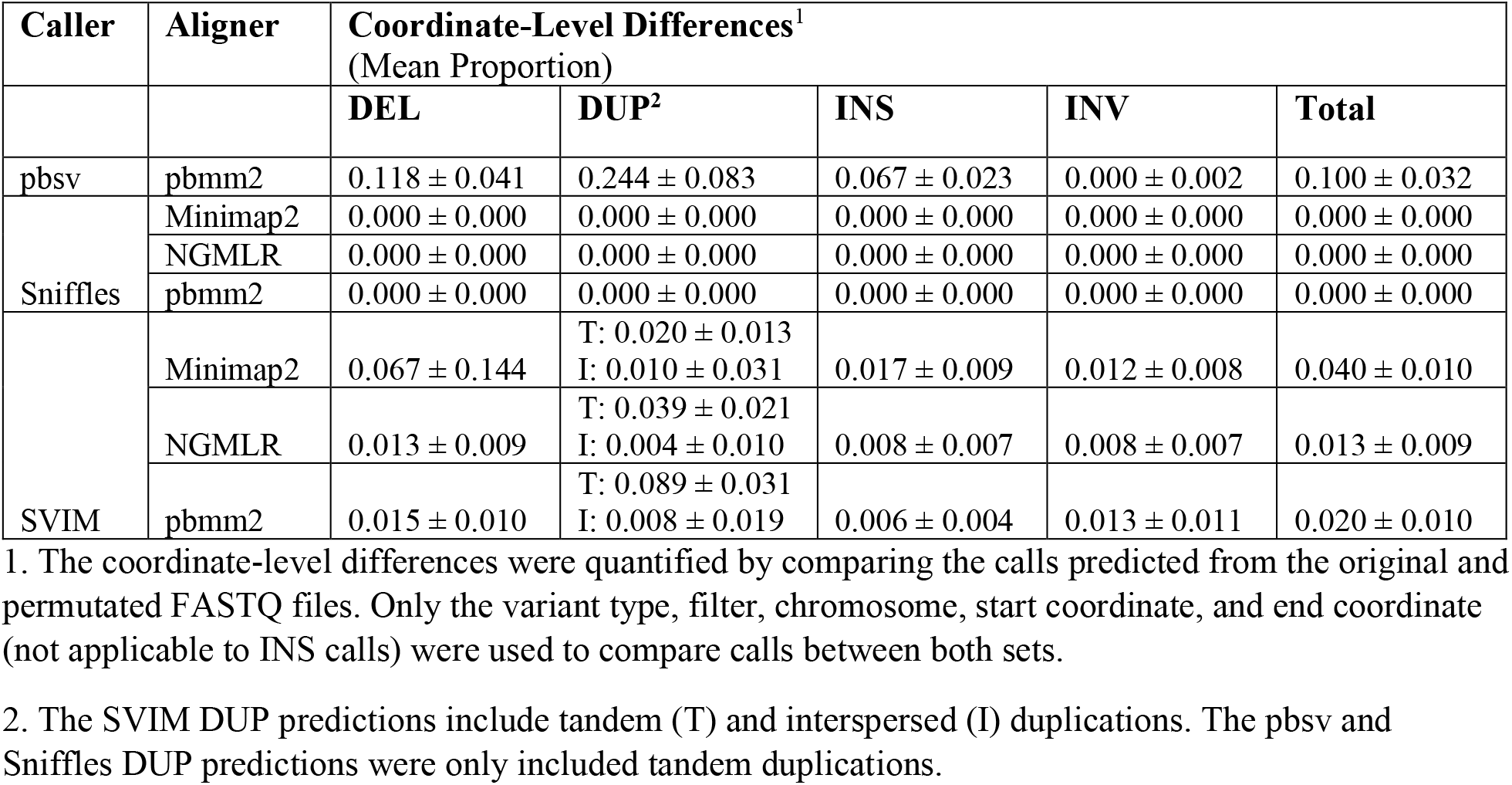
Differences in SV Calls Between the Original and Permutated Datasets Based On Genomic Coordinates.

The impact of read order varied according to the variant type in pbsv and SVIM. For pbsv, most discrepancies occurred for (0.244 ± 0.083), followed by deletions (0.118 ± 0.041), and insertions (0.067 ± 0.023). For SVIM, 0.067 ± 0.144 of the deletions predicted from the Minimap2 alignments differed between call sets, whereas fewer discrepancies were generated from NGMLR (0.013 ± 0.009) and pbmm2 (0.015 ± 0.010). The aligner also affected read order dependence for the prediction of duplications in SVIM. For tandem duplications, the differences ranged from 0.020 ± 0.013 from Minimap2 aligned data to 0.089 ± 0.031 from pbmm2. For interspersed duplications, more differences were generated from NGMLR alignments (0.004 ± 0.010) compared to Minimap2 (0.010 ± 0.031) and pbmm2 (0.008 ± 0.019).

### Sequencing Depth Affects Read Order Dependence

To evaluate the impact of sequencing depth on FASTQ read order dependence in SV calling, the *C. elegans* sequencing data were subsampled to four depths (10X, 20X, 40X, 60X). For each caller, fewer differences resulted from read order randomization of the subsampled data compared to the full depth data (Tables S1-S3). Among the callers, the differences were highest in pbsv (Figure 3), which increased from 0.005 ± 0.004 at 10X to 0.187 ± 0.046 at 60X depth. The FASTQ file read order had a smaller impact on SV calling in Sniffles and SVIM. For Sniffles, the Minimap2 aligned data at 60X depth produced the highest proportion of different calls (0.022 ± 0.008). For SVIM, the highest proportion of different calls were also produced at 60X depth (Minimap2 aligned data = 0.009 ± 0.006; NGMLR aligned data = 0.009 ± 0.007; pbmm2 aligned data = 0.009 ± 0.006).

**Figure 3.**
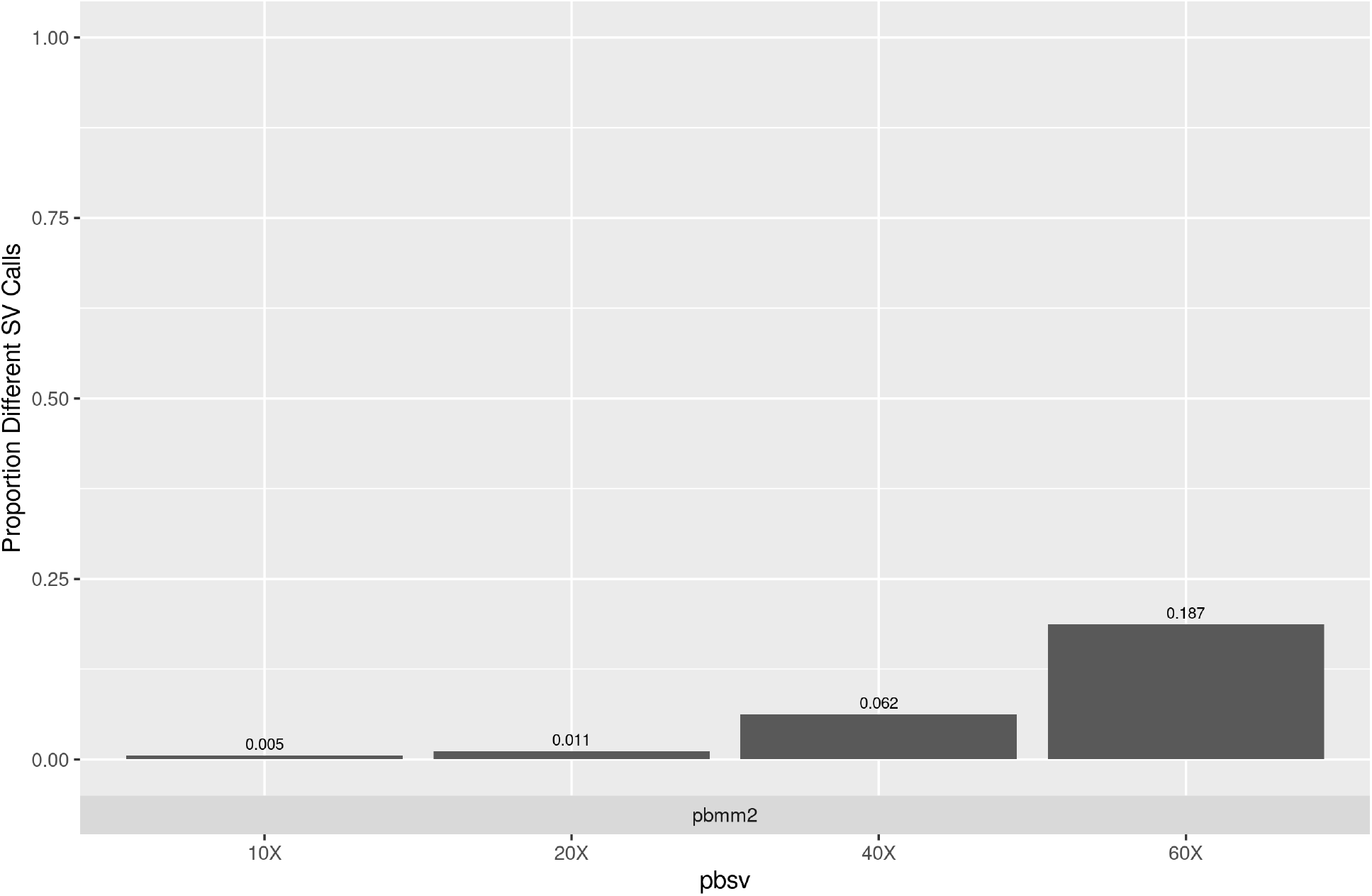
Impact of Read Depth on FASTQ Read Order Dependence in pbsv. The proportion of different calls between the SVs predicted from the original and permutated FASTQ files was used to quantify the impact of read order at four depths (10X, 20X, 40X, and 60X). All VCF file fields, except for the variant ID, were used to compare SV calls.

## Discussion

Our results demonstrate the importance of considering the impact of FASTQ read order on SV calling and further highlight the need to consider how routine intermediate methods, such as BAM sorting, can affect the results of a bioinformatics analysis. While the FASTQ file read order affected each caller, this dependence was considerably higher in pbsv. For the pbsv SVs predicted from the original and permutated FASTQ files, 60% of the total calls were unique to either call set. Conversely, only 2% of the calls differed between the predictions generated by Sniffles. The alignment method had a greater impact on SVIM compared to Sniffles, as call set differences following read order randomization ranged between 5% and 8% depending on the aligner used.

We also used subsampled data to determine if read order dependence is affected by sequencing depth. For each caller, more differences between the original and permutated data occurred using the full depth alignments to generate SV calls. However, the read order still made a considerable impact on pbsv at 40X and 60X depths. Multiple factors must be considered to estimate the optimal sequencing depth for SV calling. These include desired recall, breakpoint accuracy, sequencing platform, tool choice, and the SV types under study. Generally, this choice involves trade-offs between accuracy and cost. Although the minimum depth depends on the study requirements, past benchmarks typically range between 15X and 30X^2,10,26^. While the read order dependence may be discounted as an artifact of using exceptionally high read depths, similar or higher depths have been generated for other species, such as *Drosophila melanogaster*^27^ and *Plasmodium knowlesi*^28^.

Long-read sequencing has the potential to overcome some of the limitations of short-read approaches, but room for improvement remains for the current generation of callers. As tool developers work to improve algorithm performance, awareness of the effect of FASTQ read order on SV calling would be beneficial. This is of particular importance because only a small number of datasets with known SVs are currently available for benchmarking new tools. It needs to be emphasized that the dependence of SV callers FASTQ file read order should not be interpreted as a measure of accuracy. The analyses were chosen to quantify variance in SV calling attributable to read order. It is unclear whether SV calls that are susceptible to changes in read order are more likely to be incorrect.

Although these analyses were not chosen to quantify caller accuracy, they may assist developers in the development of more accurate tools. Benchmarks produced using the same input data for a given truth dataset, may provide misleading estimates if the tools under evaluation are dependent on read order. This is a plausible concern, as the limited availability of datasets with known SVs has resulted in most callers being benchmarked using the same data. Theoretically, if a tool is highly dependent on read order, it may be optimized for the data used to benchmark its performance. By minimizing the read order dependence, the developer may be able to provide improved estimates of the tool’s performance on other datasets.

This work is also relevant to discussions of reproducibility and replicability. Because the FASTQ file read order can affect the predicted variants, the randomness inherent in sequence order may contribute to failed replication attempts. Therefore, if future replication is a concern, we recommend that researchers sort alignment files using a tool that is insensitive to read order, such as Picard.

## Conclusions

Although many researchers are aware of the limitations of published benchmarks in bioinformatics, it is likely that differences resulting from random, arbitrary changes to the order of input data are underappreciated. Our results indicate that randomly permutating the order of reads in a FASTQ file can have a profound impact on the predicted structural variants. Seeing that the order of reads in a FASTQ file have no biological significance, we anticipate that our results will be of interest to tool developers interested in improving SV prediction. By quantifying the impact of read order, a developer may gain a better understanding of how random chance affects the relationship between the input data provided to their algorithm and the output it provides.

## Acknowledgements

We thank the University of Calgary’s Faculty of Veterinary Medicine and Research Computing Services for their investment in and maintenance of high-performance computing facilities.

