## Supplementary Tables for "The impact of FASTQ and alignment read order on structural variation calling from long-read sequencing data"

Table S1: Impact of Sequencing Depth on Read Order Dependence in pbsv

| **Depth** | **Aligner** | **DEL** | | **DUP** | | **INS** | | **INV** | | **Total** | |
|---|---|---|---|---|---|---|---|---|---|---|---|
| | | Total Calls (Mean ± StDev) | Diff. (Mean Prop.)[1] | Total Calls (Mean ± StDev) | Diff. (Mean Prop.)[1] | Total calls (Mean ± StDev) | Diff. (Mean Prop.)[1] | Total calls (Mean ± StDev) | Diff. (Mean Prop.)[1] | Total calls (Mean ± StDev) | Diff. (Mean Prop.)[1] |
| 10X | pbmm2 | 1491 ± 848 | 0.002 ± 0.003 | 186 ± 40 | 0.011 ± 0.023 | 1951 ± 944 | 0.004 ± 0.010 | 93 ± 28 | 0.003 ± 0.012 | 3721 ± 1,841 | 0.005 ± 0.004 |
| 20X | pbmm2 | 1663 ± 938 | 0.006 ± 0.015 | 265 ± 50 | 0.017 ± 0.061 | 2370 ± 1072 | 0.005 ± 0.010 | 112 ± 34 | 0.006 ± 0.015 | 4410 ± 2,067 | 0.011 ± 0.005 |
| 40X | pbmm2 | 1755 ± 1009 | 0.054 ± 0.141 | 325 ± 60 | 0.073 ± 0.135 | 2602 ± 1190 | 0.040 ± 0.116 | 122 ± 38 | 0.070 ± 0.170 | 4804 ± 2,265 | 0.062 ± 0.022 |
| 60X | pbmm2 | 1911 ± 1128 | 0.143 ± 0.205 | 378 ± 72 | 0.154 ± 0.173 | 2848 ± 1339 | 0.091 ± 0.136 | 138 ± 47 | 0.193 ± 0.262 | 5275 ± 2,553 | 0.187 ± 0.046 |

1. Proportion of different calls (mean ± standard deviation)

Table S2: Impact of Sequencing Depth on Read Order Dependence in Sniffles

| Depth | Aligner | DEL | | DUP | | INS | | INV | | Total | |
|---|---|---|---|---|---|---|---|---|---|---|---|
| | | Total Calls (Mean ± StDev) | Diff. (Mean Prop.)[1] | Total Calls (Mean ± StDev) | Diff. (Mean Prop.)[1] | Total Calls (Mean ± StDev) | Diff. (Mean Prop.)[1] | Total Calls (Mean ± StDev) | Diff. (Mean Prop.)[1] | Total Calls (Mean ± StDev) | Diff. (Mean Prop.)[1] |
| 10X | Minimap2 | 1876 ± 1071 | 0.000 ± 0.000 | 39 ± 3 | 0.003 ± 0.010 | 2647 ± 1168 | 0.003 ± 0.010 | 91 ± 35 | 0.000 ± 0.000 | 4,652 ± 2,278 | 0.001 ± 0.001 |
| 10X | NGMLR | 1703 ± 962 | 0.000 ± 0.000 | 235 ± 46 | 0.001 ± 0.003 | 2173 ± 1026 | 0.001 ± 0.003 | 209 ± 65 | 0.000 ± 0.000 | 4,320 ± 2,081 | 0.001 ± 0.001 |
| 10X | pbmm2 | 1690 ± 922 | 0.000 ± 0.000 | 71 ± 10 | 0.000 ± 0.002 | 2619 ± 1017 | 0.027 ± 0.002 | 122 ± 42 | 0.000 ± 0.000 | 4,502 ± 1,974 | 0.000 ± 0.001 |
| 20X | Minimap2 | 1998 ± 1134 | 0.000 ± 0.000 | 42 ± 13 | 0.010 ± 0.018 | 2599 ± 1201 | 0.010 ± 0.018 | 107 ± 45 | 0.000 ± 0.000 | 4,747 ± 2,385 | 0.005 ± 0.003 |
| 20X | NGMLR | 1821 ± 1024 | 0.000 ± 0.000 | 282 ± 58 | 0.010 ± 0.019 | 2250 ± 1088 | 0.010 ± 0.019 | 249 ± 81 | 0.000 ± 0.000 | 4,603 ± 2,231 | 0.005 ± 0.002 |
| 20X | pbmm2 | 1809 ± 973 | 0.000 ± 0.000 | 81 ± 11 | 0.003 ± 0.004 | 2579 ± 1066 | 0.003 ± 0.004 | 143 ± 51 | 0.000 ± 0.000 | 4,612 ± 2,083 | 0.002 ± 0.001 |
| 40X | Minimap2 | 2006 ± 1134 | 0.000 ± 0.000 | 47 ± 15 | 0.020 ± 0.021 | 2623 ± 1227 | 0.020 ± 0.021 | 124 ± 55 | 0.000 ± 0.000 | 4,800 ± 2,420 | 0.014 ± 0.007 |
| 40X | NGMLR | 1879 ± 1058 | 0.000 ± 0.000 | 323 ± 67 | 0.016 ± 0.018 | 2397 ± 1137 | 0.016 ± 0.018 | 282 ± 95 | 0.000 ± 0.000 | 4,881 ± 2,340 | 0.013 ± 0.005 |
| 40X | pbmm2 | 1868 ± 998 | 0.000 ± 0.000 | 96 ± 11 | 0.008 ± 0.011 | 2659 ± 1130 | 0.008 ± 0.011 | 167 ± 60 | 0.000 ± 0.000 | 4,791 ± 2,178 | 0.007 ± 0.004 |
| 60X | Minimap2 | 1970 ± 1109 | 0.000 ± 0.000 | 48 ± 17 | 0.030 ± 0.028 | 2708 ± 1261 | 0.030 ± 0.028 | 129 ± 59 | 0.000 ± 0.000 | 4,855 ± 2,426 | 0.022 ± 0.008 |
| 60X | NGMLR | 1898 ± 1065 | 0.000 ± 0.000 | 341 ± 69 | 0.022 ± 0.022 | 2528 ± 1190 | 0.022 ± 0.022 | 296 ± 100 | 0.000 ± 0.000 | 5,063 ± 2,400 | 0.017 ± 0.007 |
| 60X | pbmm2 | 1883 ± 1006 | 0.000 ± 0.000 | 106 ± 11 | 0.018 ± 0.026 | 2859 ± 1223 | 0.018 ± 0.026 | 178 ± 67 | 0.000 ± 0.000 | 5,026 ± 2,265 | 0.013 ± 0.007 |

1. Proportion of different calls (mean ± standard deviation)

Table S3: Impact of Sequencing Depth on Read Order Dependence in SVIM

| Depth | Aligner | DEL | | DUP: Int[2] | | DUP: TAN[3] | | INS | | INV | | Total | |
|---|---|---|---|---|---|---|---|---|---|---|---|---|---|
| | | Total Calls (Mean ± StDev) | Diff. (Mean Prop.)[1] | Total Calls (Mean ± StDev) | Diff. (Mean Prop.)[1] | Total Calls (Mean ± StDev) | Diff. (Mean Prop.)[1] | Total Calls (Mean ± StDev) | Diff. (Mean Prop.)[1] | Total Calls (Mean ± StDev) | Diff. (Mean Prop.)[1] | Total Calls (Mean ± StDev) | Diff. (Mean Prop.)[1] |
| 10X | Minimap2 | 3303 ± 1543 | 0.004 ± 0.019 | 29 ± 22 | 0.000 ± 0.000 | 558 ± 146 | 0.000 ± 0.000 | 18155 ± 3157 | 0.021 ± 0.041 | 128 ± 38 | 0.000 ± 0.000 | 22,731 ± 4,164 | 0.002 ± 0.001 |
| 10X | NGMLR | 2267 ± 1137 | 0.011 ± 0.062 | 12 ± 10 | 0.000 ± 0.000 | 1357 ± 320 | 0.002 ± 0.006 | 13933 ± 2086 | 0.011 ± 0.052 | 370 ± 84 | 0.000 ± 0.000 | 17,956 ± 3,305 | 0.003 ± 0.002 |
| 10X | pbmm2 | 2612 ± 1160 | 0.000 ± 0.001 | 50 ± 34 | 0.000 ± 0.000 | 978 ± 151 | 0.008 ± 0.020 | 18917 ± 3127 | 0.003 ± 0.009 | 212 ± 70 | 0.000 ± 0.000 | 22731 ± 4164 | 0.002 ± 0.001 |
| 20X | Minimap2 | 4363 ± 1933 | 0.010 ± 0.016 | 22 ± 17 | 0.006 ± 0.021 | 813 ± 222 | 0.000 ± 0.001 | 32934 ± 5044 | 0.014 ± 0.015 | 185 ± 65 | 0.000 ± 0.000 | 39,784 ± 5,837 | 0.003 ± 0.001 |
| 20X | NGMLR | 2679 ± 1275 | 0.001 ± 0.005 | 41 ± 32 | 0.000 ± 0.000 | 1885 ± 431 | 0.001 ± 0.002 | 25745 ± 4082 | 0.005 ± 0.007 | 517 ± 127 | 0.000 ± 0.000 | 30,877 ± 5,182 | 0.003 ± 0.002 |
| 20X | pbmm2 | 3241 ± 1291 | 0.002 ± 0.007 | 87 ± 51 | 0.000 ± 0.000 | 1438 ± 216 | 0.015 ± 0.019 | 34787 ± 4902 | 0.005 ± 0.006 | 296 ± 103 | 0.000 ± 0.000 | 39784 ± 5837 | 0.003 ± 0.001 |
| 40X | Minimap2 | 5707 ± 2347 | 0.014 ± 0.021 | 36 ± 24 | 0.013 ± 0.028 | 1154 ± 321 | 0.001 ± 0.003 | 57429 ± 8089 | 0.024 ± 0.021 | 283 ± 121 | 0.000 ± 0.002 | 66,484 ± 7,735 | 0.005 ± 0.003 |
| 40X | NGMLR | 3236 ± 1425 | 0.007 ± 0.026 | 55 ± 38 | 0.007 ± 0.015 | 2580 ± 561 | 0.003 ± 0.003 | 47652 ± 7041 | 0.010 ± 0.010 | 717 ± 199 | 0.000 ± 0.001 | 54,272 ± 8,050 | 0.005 ± 0.004 |
| 40X | pbmm2 | 4055 ± 1476 | 0.007 ± 0.010 | 117 ± 69 | 0.006 ± 0.018 | 2034 ± 325 | 0.027 ± 0.029 | 59931 ± 6858 | 0.010 ± 0.009 | 427 ± 164 | 0.000 ± 0.001 | 66484 ± 7735 | 0.005 ± 0.003 |
| 60X | Minimap2 | 6685 ± 2655 | 0.025 ± 0.020 | 47 ± 30 | 0.030 ± 0.051 | 1429 ± 404 | 0.005 ± 0.010 | 68907 ± 6266 | 0.044 ± 0.040 | 375 ± 174 | 0.007 ± 0.013 | 71,952 ± 9,020 | 0.009 ± 0.006 |
| 60X | NGMLR | 3683 ± 1536 | 0.015 ± 0.018 | 29 ± 22 | 0.049 ± 0.105 | 3103 ± 643 | 0.007 ± 0.008 | 61504 ± 7185 | 0.026 ± 0.028 | 886 ± 263 | 0.007 ± 0.023 | 69,294 ± 8,286 | 0.009 ± 0.007 |
| 60X | pbmm2 | 4658 ± 1625 | 0.012 ± 0.012 | 12 ± 10 | 0.022 ± 0.041 | 2481 ± 422 | 0.036 ± 0.034 | 64229 ± 8552 | 0.025 ± 0.024 | 536 ± 215 | 0.004 ± 0.006 | 71,952 ± 9,020 | 0.009 ± 0.006 |

1. Proportion of different calls (mean ± standard deviation)
2. Interspersed duplications
3. Tandem duplications
